# Enabled primarily controls filopodial morphology, not actin organization, in the TSM1 growth cone in *Drosophila*

**DOI:** 10.1101/2022.06.01.494414

**Authors:** Hsiao Yu Fang, Rameen Forghani, Akanni Clarke, Philip G. McQueen, Aravind Chandrasekaran, Kate M. O’Neill, Wolfgang Losert, Garegin A. Papoian, Edward Giniger

## Abstract

Ena/VASP proteins are processive actin polymerases that are required throughout animal phylogeny for many morphogenetic processes, including axon growth and guidance. Here we use live imaging of morphology and actin organization in the TSM1 axon of the Drosophila wing to dissect the mechanism of Ena action. We find that altering Ena activity has a substantial impact on filopodial morphology in this growth cone, but exerts only modest effects on actin organization. This is in contrast to the main regulator of Ena, Abl tyrosine kinase, which has profound effects on actin and only mild effects on TSM1 growth cone morphology. These data suggest that the primary role of Ena in this axon may be to link actin to morphogenetic processes of the plasma membrane, rather than regulating actin organization itself. These data also suggest that a key role of Ena, acting downstream of Abl, may be to maintain a constant filopodial organization of the growth cone, even as Abl activity varies in response to guidance cues in the environment.

**Summary statement:** We dissect the function of the actin polymerase, Enabled, in axon growth by live-imaging of actin dynamics and axon morphology of the TSM1 neuron in its native environment in vivo.

## Introduction

As an axon grows to build a neural circuit, its trajectory is directed by cues in its extracellular environment (Dickson, 2002; Tessier-Lavigne and Goodman, 1996). One of the key cytoplasmic signaling mechanisms that transduces and integrates those external signals is the Abl protein tyrosine kinase and its associated signaling network (Bradley and Koleske, 2009; Lanier and Gertler, 2000; Moresco and Koleske, 2003). Abl is a key element downstream of many of the common, conserved cell surface receptors that direct axon growth and guidance in organisms across the animal kingdom (Deinhardt et al., 2011; Forsthoefel et al., 2005; Garbe et al., 2007; Grossman et al., 2013; Hsouna et al., 2003; Yu et al., 2001). Abl is also an upstream regulator of many aspects of cytoskeletal organization, including polymerization, branching, bundling, severing, and contractility of actin networks (Bradley and Koleske, 2009; Lanier and Gertler, 2000). This makes Abl a uniquely informative tool for dissecting the molecular mechanisms by which external signals generate neuronal morphology and connectivity.

Previous studies have revealed two distinct modes of axon growth. On rigid, highly adherent substrata, the growing tip of an axon – the growth cone – often assumes a flat, lamellar morphology that employs a form of adhesive growth that links retrograde flow of intracellular actin to fixed contact points on the substratum (Lin and Forscher, 1995; Lowery and Van Vactor, 2009; Sheetz et al., 1998; Suter and Forscher, 2000). In contrast, in compliant, three-dimensional conditions like the milieu most commonly encountered in vivo, growth cones tend to be preferentially or even exclusively filopodial, and advance by a protrusive mechanism based on selection and stabilization of appropriately-oriented projections, and largely independent of substratum adhesion (Clarke et al., 2020a; Clarke et al., 2020b; Gomez and Letourneau, 1994; Sabry et al., 1991; Santos et al., 2020). This protrusive mode of growth is common across phylogeny (Leung and Holt, 2012; Murray et al., 1998; Ozel et al., 2015; Sainath and Granato, 2013; Sanchez-Soriano et al., 2010), but its mechanism has received far less attention than has the adhesive mode.

The protrusive mode of axon growth, and its regulation by Abl-dependent cytoplasmic signaling, have been studied in greatest detail by simultaneous live imaging of fluorescent markers for actin and plasma membrane in the extending TSM1 sensory axon of the developing Drosophila wing (Clarke et al., 2020a; Clarke et al., 2020b). These studies revealed, first, that the position of the protrusive, filopodial domain of the axon, the growth cone, is determined by the presence of a mass of actin in the core of the axon shaft, which provides the raw materials required for growth cone filopodia, and that forward motion of that actin mass is the driving force that, in response, induces advance of the emergent filopodial domain. Those experiments further revealed that the role of Abl kinase is to coordinate the expansions and contractions of the actin mass, introducing a spatial bias into its stochastic fluctuations that have the net effect of advancing the actin distribution down the axon. Specifically, increased Abl activity causes net expansion of the actin gel, while decreased Abl causes net condensation. Thus, a pattern of guidance cues that generates a gradient of Abl activity across a growth cone could, in principle, cause preferential expansion of actin at the leading edge of the growth cone and contraction away from the trailing edge. This would produce net advance of the actin mass over time, and therefore net advance of the morphological growth cone and growth of the axon.

While experiments to date have characterized the effect of Abl on the spatial distribution of actin in the TSM1 growth cone, it is now essential to dissect the molecular mechanism downstream of Abl by which the effects of the kinase control the distribution of actin. The best characterized, and most direct, effector linking Abl to actin dynamics is the actin polymerase, Enabled (Ena) (Gertler et al., 1995; Krause et al., 2003). Ena promotes growth of actin filaments, both as a processive polymerase that juxtaposes G-actin monomers with the barbed ends of F-actin filaments(Winkelman et al., 2014), and also by antagonizing the binding of actin filament capping proteins(Bear and Gertler, 2009; Gates et al., 2009). Ena also bundles actin filaments, in part by forming tetramers that link adjacent filaments(Bruhmann et al., 2017). Ena has profound effects on cell morphology. Overexpression of Ena stimulates formation of filopodia in many systems, though the mechanism by which it does so is complex and context dependent(Gates et al., 2007; Krause et al., 2002; Lebrand et al., 2004; Trichet et al., 2008). For example, the properties of filopodia induced by Ena alone are rather different to those produced in conjunction with formins, such as Drosophila Diaphanous(Bilancia et al., 2014; Homem and Peifer, 2009). Moreover, in some contexts Ena can promote or stabilize lamellipodia, in part by extending the actin filaments that line the leading edge of such structures(Lacayo et al., 2007; Rottner et al., 1999). The effects of Ena on morphogenesis are not determined solely by its direct effects on actin, however. Ena proteins have a conserved EVH1 domain that binds the peptide motif FPPPP (FP4), which is commonly found in adhesive structures, such as focal adhesions(Bear et al., 2000). Ena can therefore link the actin cytoskeleton to the plasma membrane. Ena is present in axons and has demonstrated effects on axon growth and guidance in vitro and in vivo(Kuzina et al., 2011; Lebrand et al., 2004; Wills et al., 1999). The molecular mechanisms underlying those effects have so far been enigmatic, however(Krause et al., 2002). First described in Drosophila, Ena has close orthologs in C. elegans (UNC-34) (Fleming et al., 2010; Sheffield et al., 2007) and in mammals (MENA, VASP (vasodilator-stimulated phosphoprotein) and EVL (Ena- and VASP-like)) that have similar properties(Gertler et al., 1996). Ena was first identified as a genetic antagonist of Abl, in that the phenotypes of *Abl* mutants can be suppressed by reducing the gene dosage of *ena*(Gertler et al., 1995), and at least some phenotypes of *Abl* mutant animals seem to be produced by mislocalization and/or hyperactivity of Ena protein(Grevengoed et al., 2003; Grevengoed et al., 2001; Kannan et al., 2014). Genetic tests show that *ena* acts downstream of *Abl*, and consistent with this, Abl regulates Ena activity, in part, by phosphorylation of conserved tyrosine residues(Comer et al., 1998). This cannot be the entire mechanism by which Abl regulates Ena, however, as an Ena derivative lacking these tyrosines still retains significant activity to perform its Abl-dependent functions(Comer et al., 1998).

Here we perform live imaging of axon morphology and actin distribution in TSM1 axons growing in their native environment of the developing Drosophila wing, both in wild type, and upon increase or suppression of Ena activity in the neuron. We find that altering Ena activity has a substantial effect on filopodial number and length, but that the growth cone is far more sensitive to reduction of Ena from its wild type level than it is to increase of Ena. This may suggest that the level of Ena in the wild type TSM1 axon is already close to saturating for its morphological functions. The effects of Ena on actin organization are quantitatively much more modest than those on filopodial morphology, though a sensitive analytical method reveals that reducing Ena activity tends to broaden the actin distribution relative to higher levels of Ena. This is consistent with our previous analysis of the effects of Abl in TSM1, but it is striking that Abl had a far more pronounced effect on actin than on morphology, opposite to our observations here of the consequences of altering Ena. Despite that, as for Abl, we find that either increasing or decreasing Ena activity causes the temporal evolution of the actin distribution to be less orderly and predictable in individual axons than what we observe in wild type. Together, these data suggest that the main role of Ena in the TSM1 growth cone may not be to regulate the actin distribution itself, but rather to modify the linkage of that actin to morphogenetic processes of the plasma membrane. Moreover, it suggests that a key function of Ena may be to buffer the downstream morphological consequences of altering Abl activity, thus maintaining the growth cone in an optimal morphology for orderly movement while still allowing the cue-directed modulation of Abl activity, and thus actin organization, that is necessary to produce guided axon growth.

## Results

We performed live imaging of the TSM 1 axon as it grows through the Drosophila wing, much as we have described previously (Clarke et al., 2020a; Clarke et al., 2020b) (Fig 1A; Suppl Fig 1). Wing imaginal discs were dissected ∼9hrs after the onset of metamorphosis (APF; <u>a</u>fter <u>p</u>uparium formation), mounted in culture media, and imaged for 90 minutes by collecting z-stacks using spinning disc confocal microscopy (interframe interval = 3 min). Membrane and actin distribution, respectively, were visualized by co-expression of CD4-td-Tomato and LifeAct-GFP, both under control of *neuralized-GAL4* (*neur-GAL4*). Axon morphology was traced in three dimensions and core growth cone parameters were quantified as described previously (Clarke et al., 2020a; Clarke et al., 2020b), including both morphological features, and actin distribution along the axon shaft (measured by radial integration of LifeAct intensity as a function of position along the axon)(Fig 1 B - D).

**Fig 1.**
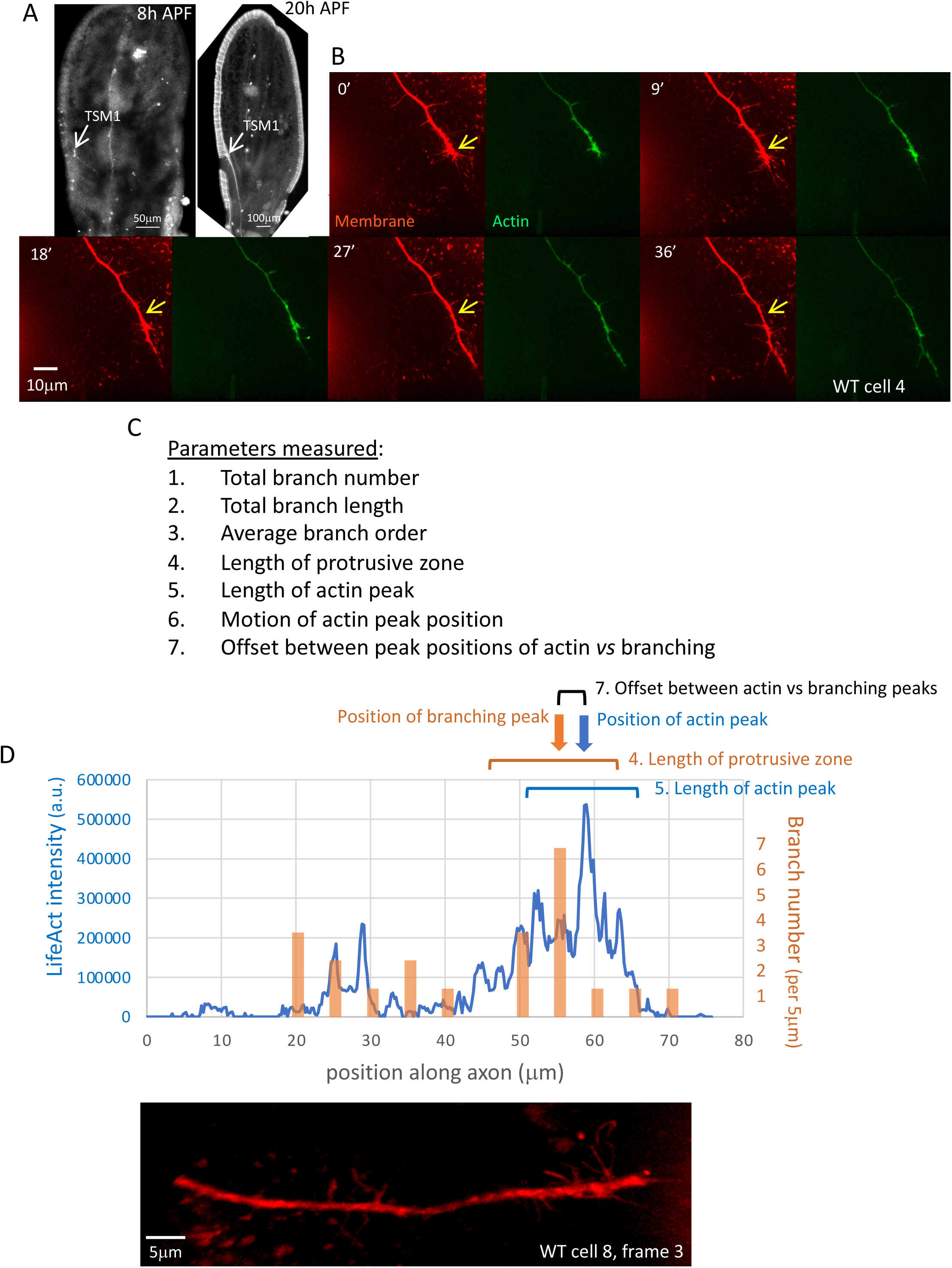
Anatomy and measured parameters of TSM1. A. Anatomy of TSM1. Z-projections of wing discs fixed and dissected at the indicated times. Neuronal membranes were labelled with anti-HRP (8hr APF image) or by expression of CD4-tdTomato (20hr APF image). Position of TSM1 cell body is indicated by white arrow; scale bar is at the bottom of each image. B. Time course of development of a typical wild type TSM1 axon, with membrane and actin, labelled, respectively, by expression of td-Tomato and LifeAct-GFP. Z-projections are shown of image stacks collected at the indicated times by spinning disc confocal microscopy. Yellow arrow in the membrane images indicates a fixed position in the disc for comparison of axon length. Scale bar is indicated in the 18’ image. C. List of parameters measured in all time points of all trajectories imaged. See Methods for details. D. For illustrative purposes, example of some of the quantified parameters from a single time point, taken from the axon whose membrane channel is shown beneath. Blue line shows the distribution of integrated LifeAct fluorescence intensity along the axon, expressed in arbitrary units (a.u.); orange bars show the number of filopodial branches arising from the axon shaft in 5 μm windows along the axon. Note that alignment with the image below is not exact as length is measured in 3-dimensional space, not in projection, and since some filopodia are not visible at this projection angle. Positions of the peaks of actin and of projections were identified using a 5μm sliding window, and are indicated with arrows. ‘Lengths’ of the peak zones of branches and of actin were calculated as the square root of the 2^nd^ moment of each distribution about its peak position, a measure corresponding essentially to ± one standard deviation (brackets). See Methods and (Clarke et al., 2020a) for detailed explanation of the utility of this definition.

Live imaging of TSM1 revealed the growth cone to be a domain of three-dimensional filopodial protrusiveness. We saw no evidence of large lamellipodial structures or obvious signs of substratum adhesion. We also found that the protrusive region of the axon, the morphological “growth cone”, contained a high local concentration of actin intensity in the axon shaft, such that the position of the window containing the peak density of protrusions correlated roughly with the position of the peak level of actin intensity (Fig 2A, B). In detail, however, we found that the peak of actin intensity tended to lead the peak of filopodial density (by 5.2 ± 0.7 μm; mean +/- SEM (median = 1.1 μm, p < 0.0001; Wilcoxon signed-rank)). Moreover, the magnitude of this offset between the positions of the actin and protrusion peaks at any given time correlated with the rate at which each one then advanced. Thus, for example, time points where actin led protrusions by a large amount were associated with greater advance of the filopodial distribution in the subsequent time step, but less advance of the actin distribution, or even its regression (Fig 2 C, D). This is consistent with the hypothesis that the offset between the actin and filopodial distributions provides the driving force for advance of the protrusive domain (Clarke et al., 2020a; Clarke et al., 2020b).

**Fig 2.**
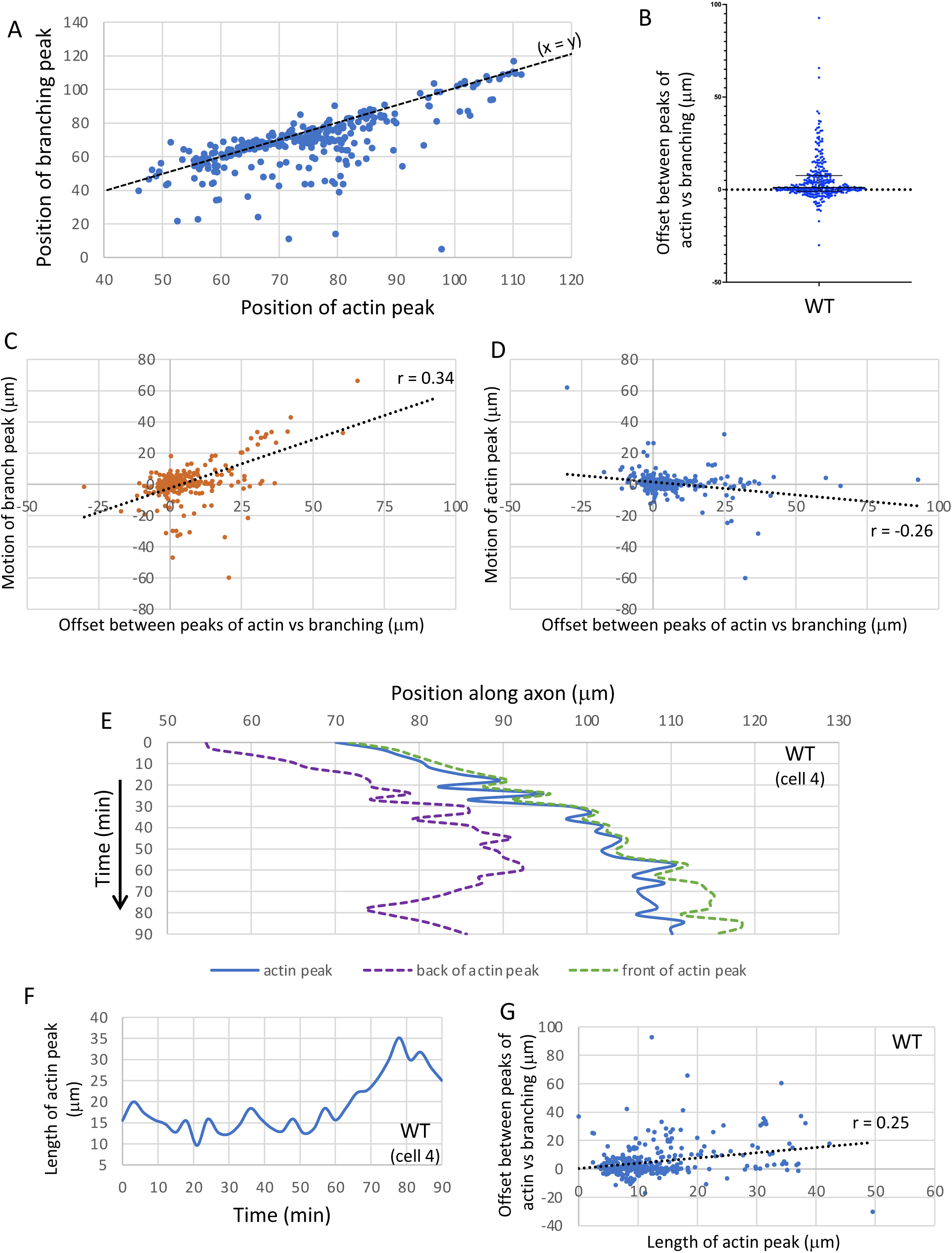
Dynamics of the growth of wild type axons. A. Scatterplot of the positions of the peak of filopodial density (“branching peak”) vs the actin peak for the entire wild type dataset. Note that many datapoints are concentrated along the line x = y (dashed black line); ie., positions of the actin and branching peaks correspond closely for many time points, however it is clear by inspection that the branch peak lags significantly behind the actin peak in a substantial number of time points – the datapoints lie well below the line. B. Tabulated offset between the positions of the peaks of actin and branching (in microns). Positive values are time points where the actin peak leads the branching peak. Median and interquartile range are shown; note that median > 0, and that the distribution is skewed heavily to larger positive than negative values. C. Scatterplot of the motion of the branch peak (in microns) in the time interval t -> t+1 vs the offset between actin and branch peaks at time t. Spearman r = 0.34; p < 0.0001. D. Scatterplot of the motion of the actin peak (in microns) in the time interval t -> t+1 vs the offset between actin and branch peaks at time t. Spearman r = -0.25; p < 0.0001. E. Graph of the positions of the actin peak (blue), and of the trailing (purple) and leading (green) ‘edges’ of the actin peak, as a function of time, in a typical wild type trajectory. Trailing and leading edge positions were calculated as, respectively, the square root of the trailing and leading partial 2^nd^ moments of the actin distribution. These are indicated as dashed lines to emphasize that they are statistical measures, not discrete axonal features. Positions are given in microns; time in minutes. Time axis increases downwards. Note the inconsistent nature of actin advance, with the peak position wiggling back and forth within the window that defines the actin mass at any given time. F. Length of the actin peak (square root of the 2^nd^ moment of the distribution) as a function of time in the same trajectory shown in E. Note the fluctuating nature of the length of the actin peak. The peak length graphed here, and used in all statistical analyses below, is based on the global second moment, whereas the positions graphed in panel 2E are calculated from the separate trailing and leading partial second moments. In general, the global 2^nd^ moment does not equal the sum of the partial 2^nd^ moments because of how these properties are calculated. The global moment is more appropriate for further statistical analyses, while the partial moments are more informative for aligning to visible features of the axon. G. Scatterplot of the offset between the positions of the actin and branching peaks vs the length of the actin peak. Spearman r= 0.25; p < 0.0001.

Detailed examination of actin distribution in the TSM1 growth cone revealed that the mass of actin undergoes constant, seemingly stochastic, fluctuations in position, but with a small, yet persistent, forward bias that produced net advance of the actin distribution over time (Fig 2E). Thus, while the position of peak actin intensity took a significant number of steps both forward and backward in any given trajectory, and these could be of roughly comparable magnitude, the net effect over the course of imaging was that the peak position of actin intensity preferentially moved forward along its trajectory. These fluctuations in peak position were also associated with fluctuations in the spatial extent of the actin peak along the length of the axon (Fig 2F), suggestive of the actin “inchworming” forward over time as it moved forward in the axon shaft. In essence, a combination of preferential forward expansion of the leading edge of the actin mass, together with preferential condensation from the rear, causes the length of the actin peak to fluctuate around a mean, but with net forward motion of the distribution midpoint. Together, these observations suggest that TSM1 growth cone advance is driven by forward-biased fluctuations of the actin distribution. This hypothesis is also consistent with the observation that the magnitude of the offset between the peaks of actin vs projection density in any given image correlated with the length of the actin peak (Fig 2G).

We next determined the effect produced on TSM1 morphology and motility when we altered the activity of Enabled (Ena) in the neuron by taking advantage of the yeast transcriptional activator, GAL4, expressed under control of regulatory sequences of the gene *neuralized* (*neur-GAL4*). Ena activity was either increased, by expressing *UAS_G_-ena(WT)*, or suppressed, by sequestering Enabled protein to mitochondria using expression of a *UAS_G_-FP4-mito* transgene. FP4-mito has an Ena binding motif, including the sequence FPPPP, linked to a mitochondrial targeting sequence. It has been validated extensively in multiple organisms and developmental contexts, and shown to provide an effective (albeit not perfect) mimic of the *ena* genetic loss-of-function condition (Bear et al., 2000; Gates et al., 2007; Kuzina et al., 2011; Lebrand et al., 2004). Its use here allows us to inactivate Ena selectively in neural tissue and bypass the lethality of a genetic null mutant.

We first found, as also observed in embryonic axon patterning (Gates et al., 2007; Wills et al., 1999), that altering Ena activity in TSM1 has relatively mild effects on overall axon patterning and on the efficiency of axon growth. We did not observe a significant frequency of defects in the final pattern of the TSM1 projection in the wing (n ≥ 30 wings for analysis of the terminal phenotype for each genotype). Moreover, the average rate of axon growth was not significantly different in the three genetic backgrounds (average growth rate = 0.20 ± 0.03 μm/min in wild type (this and all parameter values are presented in the text as mean ± SEM; see statistical methods for details of how statistical significance was calculated) vs 0.09 ± 0.08 μm/min in *UAS-FP4-mito* and 0.17 ± 0.04μm/min in *UAS-ena*; differences not significant: p=0.42; n= 10 live-imaged trajectories for wild type, 10 for *UAS-FP4-mito* and 13 for *UAS-ena* for this and all live-imaging experiments presented here)(Fig 3A). Examining the pattern of motility in greater detail revealed that the mode of growth cone movement in the altered-Ena conditions resembled that of wild type, displaying a stuttering, inconsistent pattern of advance, overlaid by stochastic fluctuation of the length of the actin mass in the growth cone, and with the actin peak tending to lead the peak of projection density (Fig 3 B-F).

**Fig 3.**
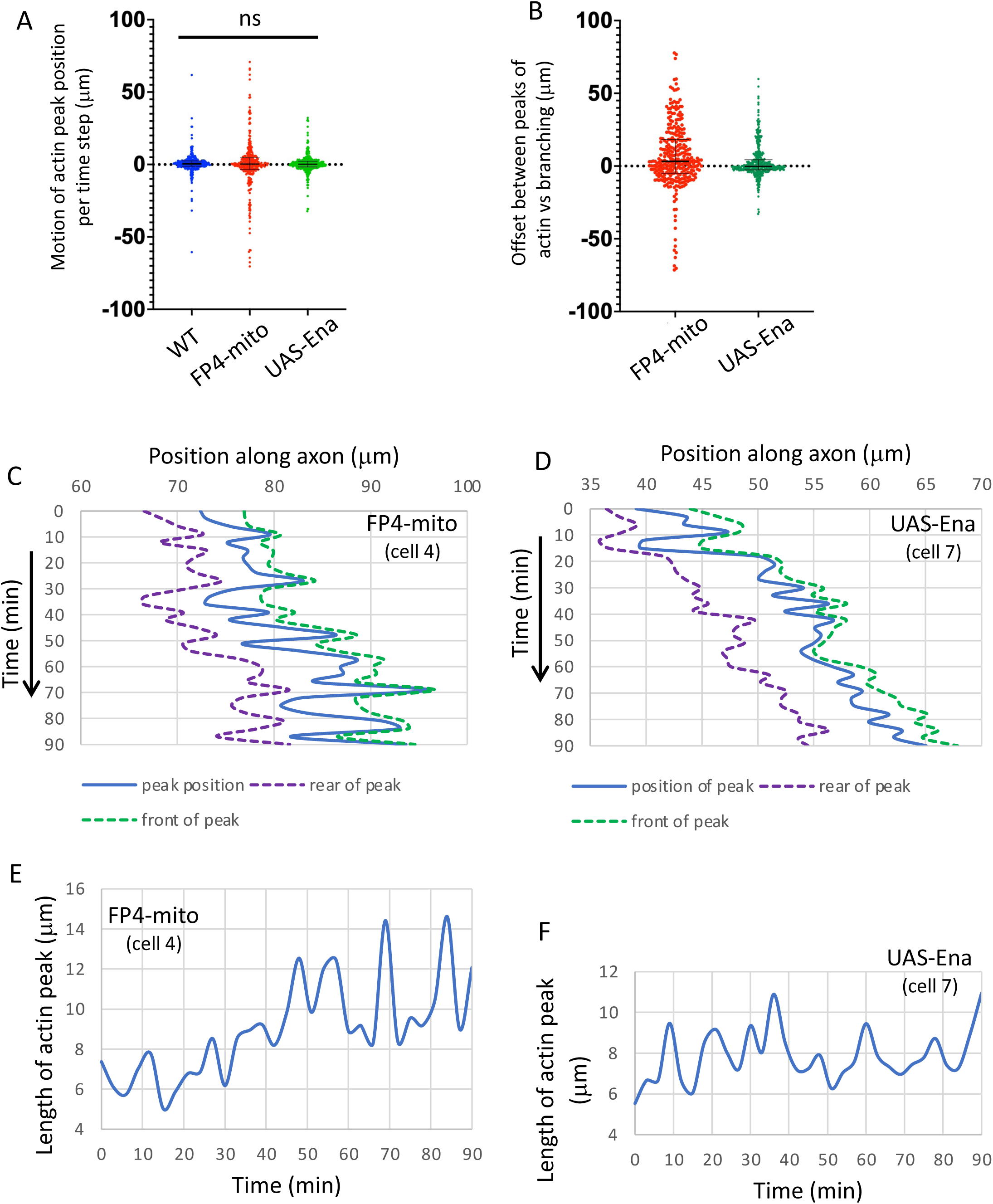
TSM1 axons with Ena loss- or gain-of-function grow similarly to wild type axons. A. Tabulated values of the motion of the actin peak between successive time points in all genotypes. Distances in microns; 3’ time steps for all trajectories. Median and interquartile ranges are shown. B. Tabulated values of the offset between the positions of the actin and branching peaks in all time points of *ena* loss of function (*UAS-FP4-mito*) and gain of function (*UAS-ena*). Median and interquartile range are shown. Median offset is significantly positive for *UAS-FP4-mito* (p<0.0001; Wilcoxon signed rank); for *UAS-ena* the median is not different from 0 by a statistically significant amount (p > 0.05), but interquartile range shows that the distribution trends toward positive values (actin peak leading branching peak). Compare with Fig 2B for the offset values in wild type. C, D. Graph of the positions of the actin peak (blue), and of the trailing (purple) and leading (green) ‘edges’ of the actin peak, as a function of time, in typical trajectories of actin loss- and gain-of-function, as described in Fig 2E. E, F. Length of the actin peak as a function of time in the same trajectories shown in C, D. Compare with wild type (Fig 2F).

We next examined the effects of Ena on the detailed pattern of actin organization and filopodial morphology in TSM1. Whereas we had found previously that altering Abl kinase activity primarily modulates actin organization, with only modest effects on growth cone morphology, we now found that the effects of manipulating Ena were opposite, primarily modifying morphology, with only modest effects on actin organization (Fig 4). First, we found that Ena activity was limiting for the accumulation of filopodial projections from the distal axon. Thus, neurons overexpressing Ena (*UAS-ena*) had 35.6±2.1 filopodia, vs 16.9±0.7 filopodia for *UAS-FP4-mito* (mean ± SEM; p < 10^-4^; ANOVA) (Fig 4A). Similarly, Ena also limited the total length of filopodial projections per axon (254.5+18.1 μm for *UAS-ena* vs 145.9±7.6 μm in *UAS-FP4-mito*; p < 10^-4^; ANOVA) (Fig 4B). For each of these parameters, comparison to wild type reveals that the neuron was far more sensitive to reduction of Ena activity than to its increase, with reduction accounting for 85% of the difference in mean filopodial number between Ena overexpression vs FP4-mito mediated suppression, and essentially 100% of the difference in total filopodial length, (filopodial number in wild type = 30.7±2.9; total filopodial length = 256.2±26.9 μm). Perhaps surprisingly, average filopodial length was not reduced upon expression of *UAS-FP4-mito*, as the decrease in total filopodial length was in proportion to the decrease in filopodial number (average filopodial length 8.9±0.4 μm in *UAS-FP4-mito* vs 8.2±0.1 in wild type; difference not significant: p=0.3; ANOVA) (Fig 4C). In contrast, the combination of increased filopodial number without a corresponding increase in total filopodial length manifested as a significant <u>decrease</u> in average filopodial length in the Ena-overexpressing condition (7.2+0.1 μm; p < 10^-4^ compared to wild type; ANOVA). Finally, in contrast to filopodial number and length, altering Ena activity did not change the complexity of filopodial projections, as the average projection order (primary, secondary, tertiary, etc.) was not altered by changes in Ena activity (1.44, 1.42, and 1.46, respectively, for wild type, *UAS-FP4-mito* and *UAS-ena*; p=0.62 across genotypes; ANOVA)(Fig 4D).

**Fig 4.**
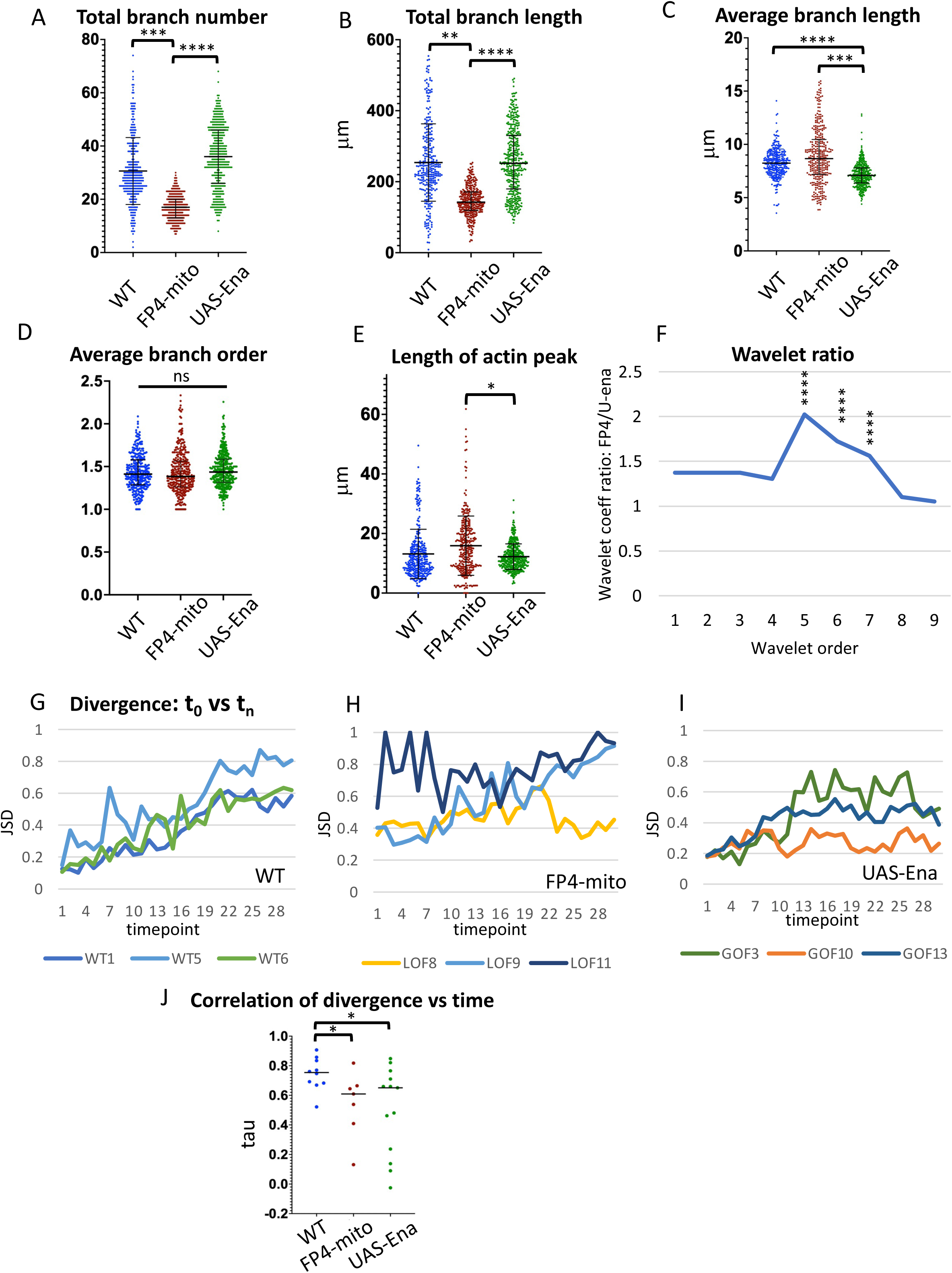
Comparison of single growth cone parameters in wild type vs altered-ena conditions. Values of indicated growth cone parameters were tabulated for all three genotypes. Statistical significance of differences is as indicated. Error bars indicate mean and SD. Statistical significance is indicated as follows for this and subsequent figures : *p < 0.05; **p < 0.01; ***p < 0.001; ****p < 0.0001. Comparisons that are not marked were not formally significant. See Methods for details of how significance was assessed. A. Total branch number B. Total branch length C. Average branch length D. Average branch order E. Length of actin peak (square root of the global 2^nd^ moment of the actin distribution). F. Plot of the ratio of the average coefficient for each spatial order of the wavelet transform for FP4-mito/UAS-ena. Wavelet analysis quantifies the spatial structure of the actin distribution, with higher-order wavelets reflecting the frequency of local actin concentrations at short length scales, and lower-order wavelets reflecting spreading of actin density at longer length scales (see text and methods, and also (Clarke et al., 2020b) for additional explanation). The maximum at wavelet order 5 indicates that expression of FP4-mito causes the actin distribution to be spread out significantly on a length scale peaking at 6.5-25.6 μm, relative to that in UAS-ena, consistent with broadening of the actin peak by reduction of Ena activity. Asterisks indicate the significance of the difference in coefficient values for the indicated order between these two genotypes. For comparison of each altered-Ena genotype to wild type, and table listing the spatial range corresponding to each wavelet order, see Suppl Fig 2. G – I. For each trajectory of each genotype, Jensen-Shannon Divergence (JSD) of the shape of the actin distribution was calculated between the first time point vs each subsequent time point. Divergence can vary from 0 (identical distributions) to 1 (unrelated distributions). For clarity, plots are shown here for only three illustrative trajectories of each genotype. For plots of JSD vs time for all trajectories, see Suppl Fig 3. J. Tabulation of the Kendall tau value of the correlation of JSD vs time for all trajectories of each genotype. Black bar indicates median value; asterisks indicate significance of genotype differences.

We next quantified the effect of Ena on parameters of actin organization. One of the key actin features of the growth cone shown previously to be regulated by Abl is the length of the actin peak. Here we saw a small shift in the expected direction, with suppression of Ena activity causing expansion of the actin peak relative to Ena overexpression (14.1μm in *UAS-FP4-mito* vs 11.9μm in *UAS-Ena*; 95% confidence intervals 11.8-16.5 vs 10.7-13.2, respectively; Fig 4E) consistent with the expansion of the actin peak seen upon overexpression of Abl, the Ena antagonist. As we found for the filopodial parameters, the majority of the difference between the two altered Ena conditions derived from the effect of Ena suppression (WT = 12.2μm; 95% CI 8.8-16.0). Quantitatively, however, the effect of Ena on the length of the actin peak was rather small in magnitude. Therefore, to confirm this observation, we performed a more sensitive analysis of the shape of the actin distribution, using wavelet analysis. The wavelet transform reveals the spatial scales of correlations in the distribution of actin along the axon. As such, it is a sensitive indicator of clustering or separation of actin density along the axon (see (Clarke et al., 2020b) for a more detailed explanation). In essence, increased values for the coefficients of higher-order wavelets arise from concentrations of actin signal at short spatial scales (small actin clumps), while lower-order wavelets reflect spreading of actin at larger spatial scales. Here we find that plotting the ratio of (wavelet amplitude)^2^ for (*UAS-FP4-mito/UAS-ena*) vs wavelet order reveals that reducing ena activity leads to a significant enhancement of the contribution of a narrow range of wavelets, peaking at 5^th^ order (p<10^-4^; ANOVA), corresponding to separation of actin density at a spatial scale corresponding to ∼6.5 – 25.6 μm (Fig 4F; Suppl Fig 2). Stated otherwise, the wavelet analysis shows that reducing ena activity causes a spread of the actin distribution at this multi-micron spatial scale, consistent with the increased length of the actin peak observed by direct measurement as Ena activity is decreased.

The other major feature of actin shown previously to be regulated by Abl in TSM1 is its degree of organization, for example, the extent to which the shape of the actin distribution at one time point predicts the shape of that distribution at a subsequent time point. We quantify this feature with a property termed the Jensen-Shannon divergence, which varies from 0 if two distributions are identical, to 1 if two distributions are entirely unrelated. For each of the wild type trajectories we analyzed, we found that the divergence between the actin distribution at time t=0 vs time t=1 is relatively small, but that the divergence increases systematically as the t=0 actin distribution is compared to later and later time points of the same trajectory (Fig 4G). In contrast, if we deregulate Ena, either by suppression with FP4-mito or by overexpression, that predictability is degraded significantly (Fig 4 H-J; Suppl Fig 3). There are still trajectories in the altered Ena conditions that show consistent increase in divergence with time, but there are also trajectories where divergence is uniformly high, or changes with time in unpredictable ways. Thus, we see that the dynamic reliability of the evolution of the actin pattern is disrupted upon deregulation of Ena.

We know from our previous analysis of Abl that single growth cone parameters with small individual responses to perturbation can nonetheless contribute to robust consequences because of consistent correlations among some growth cone features. We therefore expanded our analysis of Ena by querying the pairwise correlations of growth cone parameters, as well as examining the global effects of the whole set of growth cone features in an unbiased principal components analysis (PCA).

Correlations between individual pairs of growth cone parameters that were found to be significant across all three genetic conditions identified core features of a well-formed growth cone and of effective growth cone advance (Fig 5A; Suppl Fig 4). Thus, for example, projection number, total projection length, and the projection branching complexity (average branch order), showed significant three-way correlation in all three genotypes, suggesting that this nexus reflects a consistent feature of growth cone cell biology (Fig 5B). Consistent correlation of these three features was also observed in our previous study of TSM1 (Clarke et al., 2020a). Moreover, as discussed above for wild type, the offset between the actin and projection peaks also showed significant correlation with the length of the actin mass in both altered-Ena genotypes, as well as negative correlation between the magnitude of that offset in a given time point and the degree of advance of the actin in the following interval, both suggesting a stepwise, inchworming mode of axon growth. Also consistent with this, both altered-Ena conditions recapitulate the positive correlation of the offset between the positions of the branching and actin peaks in any given image vs advance of the filopodial distribution in the time step that followed (Fig 5 C, D), which was shown above to be a consistent feature of wild type TSM1 axon growth (Fig 2C).

**Fig 5.**
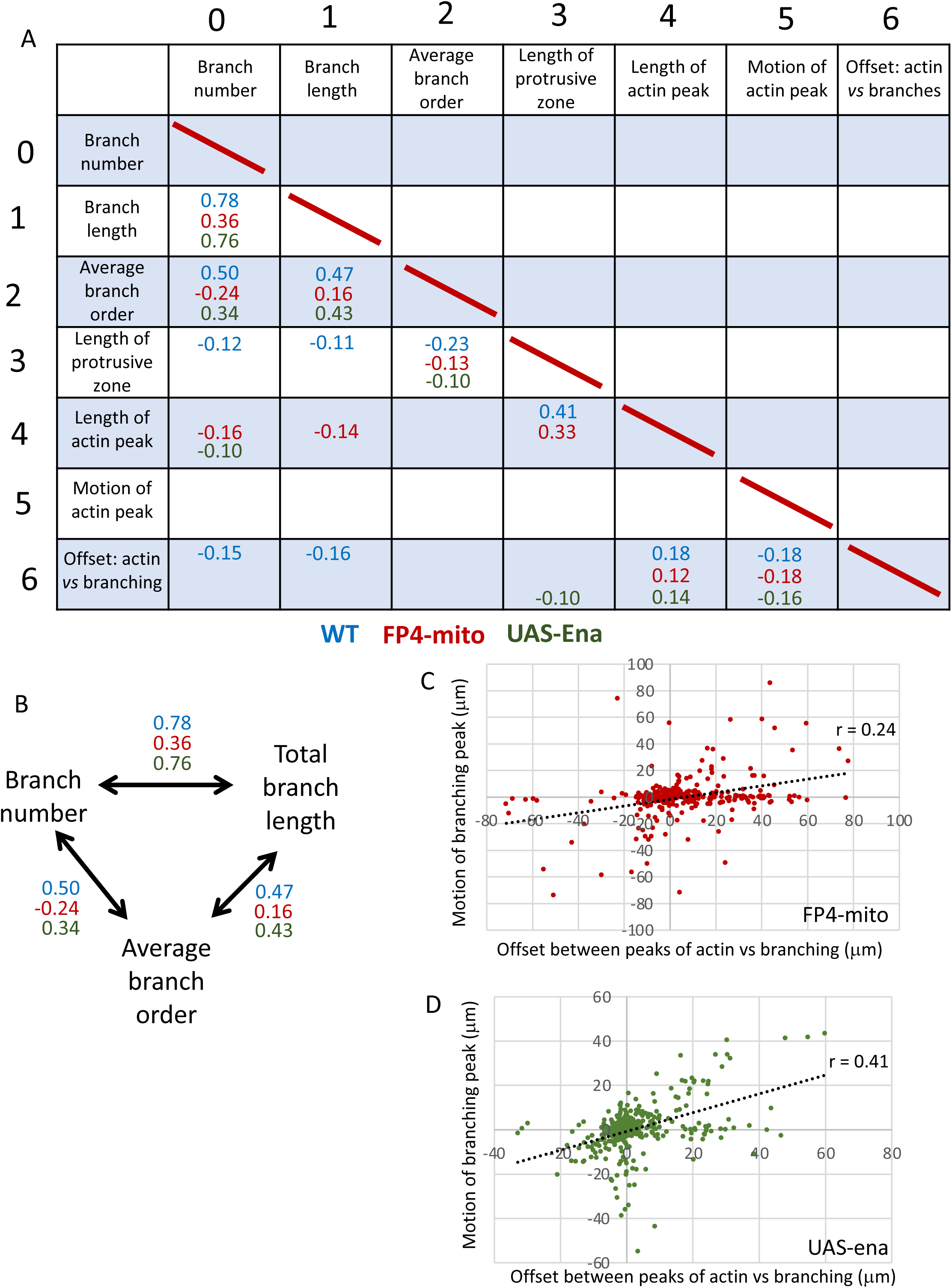
Pairwise correlations of parameters among time points of each ena genotype. A. Table of pairwise Kendall tau correlation values of the seven parameters reported for each time point. Correlation significance was calculated by the Benjamini-Hochberg method. Tau value is shown for all correlations with a false discovery rate (FDR) < 5%. Tau values in blue – wild type; red – *UAS-FP4-mito* (Ena loss-of-function); green – *UAS-ena* (Ena gain-of-function). For table of all tau values and associated p-values, see Suppl Figs 4A and B, respectively. B. Branch number, length and order were significantly correlated in all pairwise combinations in all three genotypes. Tau values are shown. C. Scatterplot of motion of the peak of branch density in a given time step vs offset between the actin and branching peaks at the beginning of that time step for *UAS-FP4-mito*. Compare to wild type (Fig 2C). Spearman r = 0.24; p < 0.0001. D. Scatterplot of motion of the peak of branch density in a given time step vs offset between the actin and branching peaks at the beginning of that time step for *UAS-ena*. Compare to wild type (Fig 2C). Spearman r = 0.41; p < 0.0001.

Unbiased global analysis of the interactions among growth cone parameters by PCA yielded additional insight into axon structure and dynamics, and how they are modulated by Ena (Fig 6A-F). Our previous study of TSM1 revealed a categorical separation of wild type growth cones into two distinct morphological classes, one with a simpler branching structure and the other more complex. The current wild type dataset reproduces that effect, revealing a bimodal distribution along PC1 (dominated by morphological features: branch number, total branch length and average branch order) with the time points from three of the wild type trajectories having PC1 values nearly exclusively less than -0.5 and the time points from the other seven trajectories almost entirely above that value (Fig 6G, H). Suppression of Ena activity by expression of FP4-mito shifted the distribution completely to the simpler morph (higher values of PC1; Fig 6D, I). In contrast, upon overexpression of Ena, while the mean value of PC1 did not change by a statistically significant amount, the distribution essentially collapsed to erase the categorical distinction between the two morphs (Fig 6J; Suppl Fig 5A). That is, rather than being obviously bimodal, the distribution now peaked in the interval between the two wild type morphological forms, with several trajectories straddling the boundary, eliminating any clear categorical distinction between morphs. We also examined PC2, which was dominated by the length of the actin distribution, and to a lesser degree by the closely-correlated length of the filopodial distribution. The three genotypes showed no discernable difference in the means of the distributions of PC2 values (Suppl Fig 5B), reinforcing the interpretation that Ena has limited effects on the actin distribution itself, in contrast to its strong effects on morphological features of TSM1.

**Fig 6.**
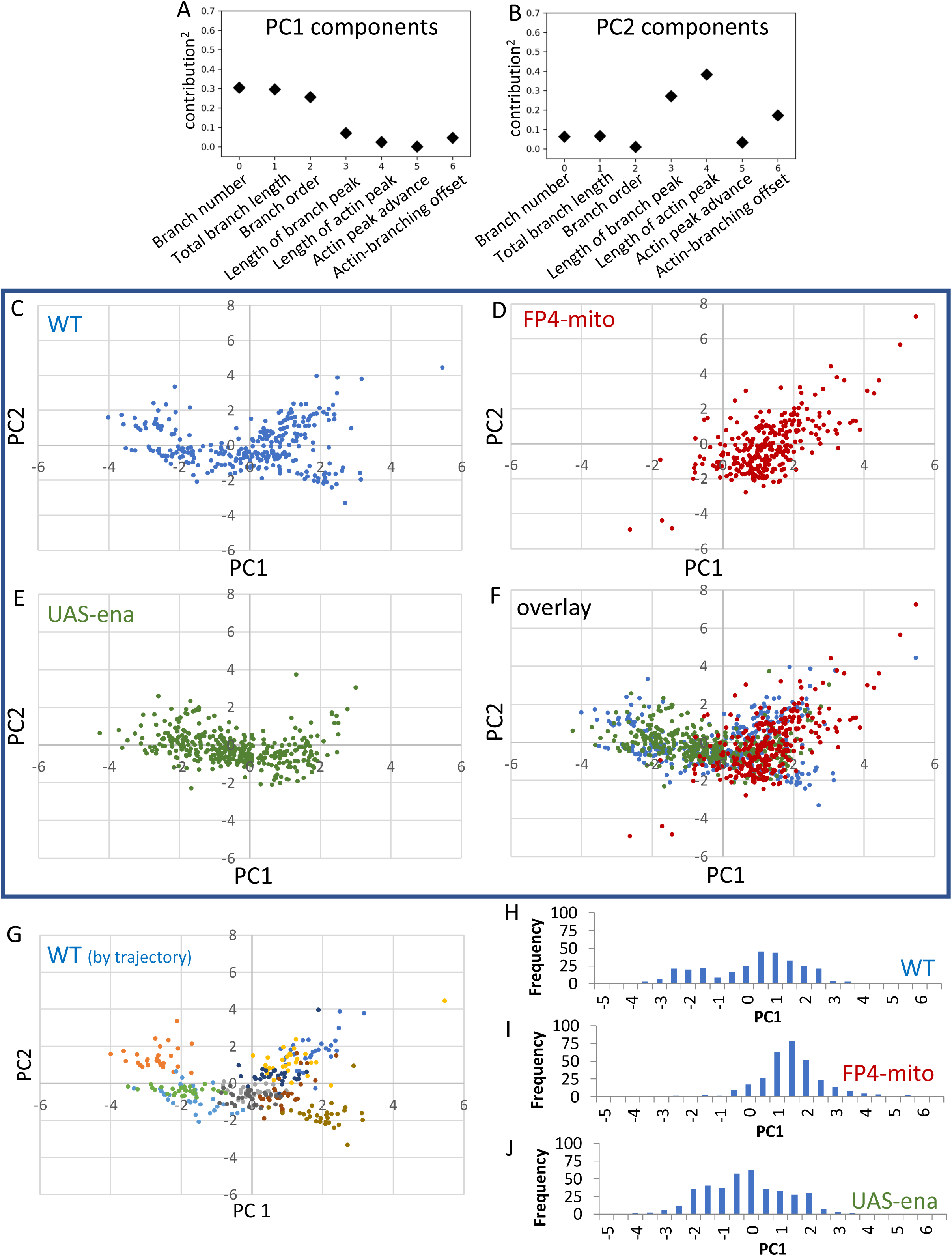
Principle component analysis of all parameters for each ena genotype. A, B. Fractional contribution to each of the first two principle components is shown for each measured parameter. (Contribution)^2^ is shown to facilitate comparison. C – F. PCA was performed on wild type using all 7 parameters, and data from FP4-mito, and *UAS-ena* datasets were then projected into the wild type PCA space. The plane corresponding to PC 1 and 2 is shown. C – E show the three genotypes individually; D shows the overlay of all 3 genotypes. G. PCA of wild type is shown with each imaged cell colored separately. Note separation of trajectories into a cluster of three cells with PC1 less than approximately -0.5, and a second cluster with PC1 greater than that value. H – J. Histogram of PC1 values for the three *ena* genotypes, as indicated. Note the bimodal distribution of PC1 in wild type; coincidence of PC1 values of FP4-mito with the higher-value wild type cluster, and PC1 values of *UAS-ena* concentrated roughly around the value of the minimum in PC1 values of wild type.

## Discussion

Here we have investigated the molecular mechanism of axon growth and guidance by using live imaging of the TSM1 axon in the developing Drosophila wing to quantify the effects of the processive actin polymerase, Ena, and to compare it with the effects of the Ena regulator, Abl tyrosine kinase. We find that Ena has a significant effect on the number of filopodial projections in the wild type growth cone, but much less effect on their length, or on where they form. We also observe an asymmetry in the effects of Ena, with suppression of Ena activity producing a large effect on growth cone morphology but increase of Ena having little effect. This may suggest that the wild type level of Ena in this growth cone is close to saturating for its morphological functions. In contrast to its substantial effects on growth cone morphology, we find that the effect of Ena on actin organization in this growth cone is quite modest. The actin mass at the heart of the growth cone undergoes slight expansion with decrease of Ena activity, but the effect size is not large. Alteration of Ena level also impairs the orderly evolution of the growth cone actin distribution over time. The limited sensitivity of the growth cone actin distribution to the level of Ena, as opposed to the strong effect of Ena on morphological features, is in striking contrast to the effect of Abl, which has a profound impact on actin organization but only modest effects on overall growth cone morphology. These observations suggest that Ena may be more important for the linkage of actin to the membrane and downstream morphogenetic processes of this axon than it is for the distribution of the actin itself. Taken together, our data also suggest that the balance of Ena with other factors regulated by Abl may serve to buffer the effects of Abl on filopodial patterning, thus maintaining an optimal growth cone morphology for orderly axon growth while still allowing Abl to act as a rheostat to vary actin distribution.

Live imaging of morphology and actin organization in a developing pioneer axon of the Drosophila wing, TSM1, recently led us to propose a novel and unexpected model for axon growth and guidance in vivo. We found that the distal part of the axon shaft contains a region with a high local concentration of actin. This actin mass undergoes constant, stochastic, fluctuations in size, but with a small forward bias that produces net advance of the actin mass over time. Since actin and associated factors are essential for building and maintaining filopodia, advance of the actin mass leads, in turn, to advance of an emergent domain of high filopodial density that is the morphological feature we recognize as “the growth cone”. We also showed that Abl tyrosine kinase is a key regulator of actin organization and dynamics in the growth cone, and thus, indirectly, of growth cone morphology and motility. In the current work, we have extended our analysis to determine the role of a core Abl effector in TSM1, the processive actin polymerase, Ena.

Consistent with data from other systems(Krause et al., 2003), we found that Ena has a significant impact on the morphology of the TSM1 growth cone in vivo. Varying Ena had a significant effect on filopodial number, with fewer filopodia present under conditions with low Ena activity than with high Ena. Surprisingly, however, nearly all of that effect (85%) was due to the effect observed upon suppressing Ena; the effect of increasing Ena (relative to wild type) was far more muted. Similarly, varying Ena activity revealed a shift to less total filopodial length with lower levels of active Ena, again with the overall effect dominated by the consequences of Ena suppression. The limited effect of Ena overexpression was unexpected, and may suggest that Ena is present in the wild type TSM1 at a level that is already nearly saturating for Ena function. A significant effect of Ena overexpression was observed, however, upon measuring average filopodial length: the combination of increased filopodial number in *UAS-Ena*, relative to wild type, without a corresponding increase in total length, manifested as a significant decrease in average filopodial length (though, curiously, suppression of Ena in this axon did not cause a decrease in average filopodial length, unlike many other systems where Ena has been investigated(Gates et al., 2007; Lebrand et al., 2004; Vasioukhin et al., 2000)). Despite these Ena-dependent changes in growth cone morphology, however, altering Ena activity did not produce overt defects in the terminal phenotype of TSM1. This is reminiscent of axon patterning in Drosophila embryonic development, where *ena* gain or loss of function also produce mutant phenotypes only in a limited set of developmental contexts(Wills et al., 1999), and also of development of retinal axons in Xenopus where Ena is important for arborization of axons, but not for their targeting (Dwivedy et al., 2007).

In contrast to these strong effects of Ena on filopodial organization, it was surprising to find that Ena has only rather subtle effects on actin organization in TSM1. A decreasing level of available Ena is associated with an increasing overall length of the actin mass in the growth cone, as well as spreading of the actin within that mass, as assayed by wavelet analysis, consistent with the expected complementarity of Ena to the effect of altering Abl, but in the case of Ena the effects are quantitatively rather small. This is different from the findings in our earlier analysis of Abl, whose major effect on TSM1 is in regulation of actin organization, with effects of Abl on morphology being much more limited. Consistent with the modest effects of Ena on actin organization in the current study, altering Ena activity did not significantly alter the average rate of advance of the growth cone actin mass. Moreover, the overall pattern of growth cone dynamics remained the same regardless of Ena activity, with longitudinal expansion of the actin mass driving instantaneous advance of the actin peak, and the resulting offset between actin mass and filopodial density driving subsequent advance of the filopodial peak.

The combination of effects we observe for Ena in TSM1 in some ways match those seen in published analyses of Ena in other systems, but in other ways they were rather unexpected. In contrast to our data here for TSM1, in other systems, Ena has been shown to have substantial effects on actin organization, and Ena overexpression has been shown to induce robust extension of cellular projections. We note, however, that those studies have in general investigated Ena action in cellular contexts dominated by substratum adhesion: epithelia(Gates et al., 2007; Vasioukhin et al., 2000), adherent cells (Bear et al., 2000; Lacayo et al., 2007), and axons growing in a lamellar fashion on rigid supports(Gupton and Gertler, 2010; Lebrand et al., 2004). In all of these cases, it may be, for example, that interaction of Ena with adhesive structures plays a critical role, feeding back on actin organization in response to morphological inputs. In TSM1, in contrast, our observations do not provide obvious suggestion of a significant adhesive contribution, similar to the limited role of adhesion in studies from other labs investigating motion in compliant, three-dimensional media, both for growth cones and for motile cells(Lammermann et al., 2008; Santos et al., 2020). Indeed, the notion suggested above that a key role of Ena in TSM1 may lie in the linkage of actin to the plasma membrane, rather than in direct effects on actin structure, would agree well with an observation that the protein has generally stronger effects in adhesion-limited cellular contexts(Sheffield et al., 2007; Vasioukhin et al., 2000) than it does in TSM1.

The experimental observations made here showing that Ena has relatively modest effects on actin organization agree well with our recent results from computational simulations of actin networks(Chandrasekaran et al., 2021). There, we found that the effects on actin from changing Ena activity were manifested most strongly on fine details of network organization at very short range (sub-micron) length scales, beyond the effective resolution of our microscopy. In the simulations, modulation of more robust actin nucleators, such as Arp2/3, and contractile elements, such as Myosin 2, were required to produce large, mesoscale (multi-micron-level) effects on actin distribution like those that we observe here in the living wing disc, and that we have shown to underlie the mechanism of axon growth and guidance. This suggests that aspects of signaling downstream of Abl that are distinct from its regulation of Ena are likely to play the key role in regulating large-scale actin organization in the growth cone. A strong candidate is Abl-dependent activation of the Rac GEF, Trio, with consequent stimulation of a Rac/WAVE pathway, which we have shown to occur in parallel to the Abl-Ena interaction (Kannan and Giniger, 2017; Kannan et al., 2017), and which would be predicted to stimulate the branching actin nucleator, Arp2/3. Moreover, it has been shown that Abl regulates the activity of Myosin 2 (Dudek et al., 2010), another key regulator of the mesoscopic organization of non-polarized actin assemblies in our simulations. Our computational analysis also suggested a simple mechanistic explanation for how nanometer-scale changes to actin filament length produce multi-micron scale changes in the overall distribution of actin density by modifying the connectivity (percolation) of the actomyosin network (Chandrasekaran et al., 2021).

While the overt effects of Ena on actin distribution that we observe here in TSM1 are relatively subtle, they are evidently significant physiologically. In particular, just as we found previously that either increased or decreased activity of Abl induces disorganization of the actin distribution, we show here that experimental manipulations that increase or decrease Ena activity, over-riding its normal regulation, disrupt the reliable evolution of the actin pattern in the growth cone over time. This is consistent with the idea that modulation of Ena activity may be important to the mechanism by which Abl ensures that transformations of the actin distribution occur in an orderly fashion in the advancing growth cone.

The data reported here show that altering Ena activity, particularly reducing Ena activity, produces strong effects on filopodial pattern in TSM1, in contrast to Abl, which has at most a mild effect on TSM1 filopodial morphology. This is curious, however. If Abl regulates Ena, and Ena strongly modifies morphology, why doesn’t altering Abl have a stronger effect on TSM1 filopodial morphology? Our earlier studies of Abl signaling may hint at an explanation. We have shown previously that there are at least two opposing signals downstream of Abl, suppression of Ena, but also activation of Trio/Rac/WAVE/Arp2-3, and we have speculated that this pattern of antagonistic regulation of its two key effectors may be critical to Abl function(Kannan and Giniger, 2017; Kannan et al., 2017). Activation of Ena promotes filopodial development, as discussed above, but so does activation of Arp2-3. It is thought that Arp2-3 promotes formation of sub-membranous, branched actin networks that nucleate the parallel actin filaments that extrude filopodia(Biyasheva et al., 2004; Korobova and Svitkina, 2008), and indeed, experimental manipulation of Arp2-3 activity has verified that activation of this protein complex enhances filopodial number in Drosophila growth cones (Sanchez-Soriano et al., 2010). Therefore, it seems plausible that the antagonistic regulation of Ena vs Arp2-3 by Abl may have the net effect of keeping the local propensity for filopodial extension in the growth cone roughly constant, even as Abl activity changes. Stated otherwise, by this model, a key function behind the complementary regulation of Ena vs Rac/WAVE/Arp2-3 may be to buffer the effects of Abl on filopodial morphogenesis, maintaining the growth cone in an optimal morphological state for continued growth, while leaving Abl activity free to be an adjustable rheostat that can be used to modulate actin organization in response to external cues, tuned to produce the directed expansion and contraction of the mesoscale actin network that is the engine for axon growth and guidance.

## Experimental Methods

### Drosophila stocks

Drosophila stocks *neur-GAL4[A101]* (BDSC 6393), *UAS-LifeAct-eGFP* (BDSC 35544) and *UAS-CD4-td-Tomato* (BDSC 35837) were obtained from the Bloomington Drosophila Stock Center (Bloomington, IN). *UAS-ena* (untagged) and *UAS-FP4-mito-GFP* were obtained from Julie Gates (Bucknell University) and Mark Peifer (UNC-Chapel Hill). Note that under our conditions of imaging (low intensity of the 488nm laser to limit photodamage and low GFP detector gain to prevent saturation of axonal LifeAct-eGFP signal; see below), GFP fluorescence was not detectable in the axon upon expression of *FP4-mito-eGFP*. Flies were raised on standard cornmeal/molasses food (KD Medical).

### Microscopy and antibody staining

Fixed samples were used only to generate the anatomical reference images of Fig 1A. To prepare the early-prepupal image, white prepupae (WPP) were collected, aged 8 hours at 25°, then dissected and fixed for 25’ in PBS containing 4% formaldehyde and 0.1% glutaraldehyde. Wings were then washed in PBS, transferred to PBS + 0.3% Triton X-100 (EM Sciences, Hatfield, PA), blocked, incubated for 90’ with TRITC-anti-HRP (Jackson ImmunoResearch, West Grove, PA; cat# 323-025-021; dilution 1:100), washed, and mounted in Prolong Gold (ThermoFisher Scientific). To prepare the mid-stage (pupal) image, WPP expressing *CD4-td-Tomato* under control of *neur-GAL4* were collected and aged 20hr at 25°. Pupae were removed from the pupal case and fixed in PBS containing 4% formaldehyde for 25’, RT. After washing, fixed pupae were dehydrated in 100% ethanol and stored in ethanol for at least 24 hrs at 4°. Pupae were then rehydrated in PBS + 0.3% Triton, wing discs were dissected and mounted in Vectashield (Vector Laboratories, Newark, CA) . Widefield microscopy was performed with a Zeiss AxioImager Z1 microscope, and image stacks were deconvoluted and processed in Zen.

### Live imaging

Live imaging was performed by a modification of the method described in Clarke, et al (Clarke et al., 2020a). WPP of the appropriate genotype were collected and aged 8hrs at 25°. Wing discs were dissected in fresh culture media (Schneider’s *Drosophila* media (Life Technologies, Frederick MD) containing 10% fetal bovine serum (Gibco)). Wing discs were transferred to a drop of culture medium (∼15μl) in the middle of an 18 x 18mm #1.5 coverslip and mounted by the method of Rusan and coworkers ((Lerit et al., 2014); see also Suppl Fig 1). In brief, discs were transferred in a minimum volume of culture medium using a pipet tip that had been treated with Sigmacote (Sigma-Aldrich, St. Louis, MO) and pre-blocked by triturating contents of the pupal abdomen. Small (∼10-15μl) drops of #700 halocarbon oil (Sigma-Aldrich) were placed at the corners of the coverslip and it was stuck to the underside of a gas-permeable Lumox 35 culture dish (Sarstedt, Numbrecht, Germany), which was then inverted. Media and oil were allowed to spread, and a kimwipe was used to wick away media and oil until wings were physically restrained but not crushed. Additional oil was used as needed to seal the edges of the coverslip. Up to 5 discs were mounted per imaging chamber. Imaging was performed on an inverted microscope, with imaging chamber right-side up and filled with ∼3 ml culture media (to avoid reflection at the surface of the dish).

Imaging was performed with a Zeiss AxioObserver Z1 spinning disc confocal microscope with a 25° temperature-controlled stage. Z-stacks were taken at 0.8 μm spacing with a 63x/1.2 NA water immersion lens. Typically, two discs were imaged at once, using the multipoint feature of the software. Imaging runs were 90 minutes with 3’ between initiation of successive frames. Images were not deconvoluted as previous experiments showed that deconvolution corrupts the information content of the images (Clarke et al., 2020a).

### Segmentation of images and quantification of growth cone parameters

Tracing of axons and quantification of LifeAct intensity were performed precisely as described in (Clarke, et al. 2020a). In brief, three-dimensional tracing in Imaris (Bitplane, version 8 or 9, Oxford Instruments, Abingdon, England) was first performed of just the axon shaft and converted to Nikon ICS format. This was imported into MIPAV (NIH, Bethesda, MD), which generated an SWC format description of the axon (plug-in: *Drosophila creates SWC*), and then calculated the LifeAct intensity as a function of position along the axon by summing signal intensity in sequential frustums centered on the axon shaft (plug-in: *3D SWC stats*). Complete tracing of all projections was then performed in Imaris, and again converted to ICS format and imported to MIPAV for preparation of an SWC file. During tracing, care was taken to begin the trace at a specific position of the proximal axon that could be identified consistently in all frames of the trajectory.

Parameters describing features of morphology and actin distribution were calculated as described previously (Clarke et al., 2020a). Custom Python scripts were written to calculate the desired parameters for each image from the SWC file of projections and from the spreadsheet of actin intensity as a function of position in the axon shaft. Parameters are listed in Fig 1C. These include the total number of projections from the axon shaft, total length and average length of projections, and average projection order. To calculate projection density along the axon, higher order projections were assigned to the position of their parent primary projection. The position of maximum projection density, and of maximum actin intensity, were identified separately using 5μm sliding windows (advanced in 1μm steps). It was shown previously that results are robust to the choice of window length (1-10μm; (Clarke et al., 2020a)). As previously, the “length” of the protrusive zone of the axon was calculated as the square root of the second moment of the distribution of filopodial density about the position of the window with the maximum value, and the “length” of the region of elevated actin concentration was similarly calculated as the square root of the second moment of the actin intensity about the position of the maximum window. The square root of the second moment is essentially analogous to one standard deviation, and was previously found empirically to a be a useful measure of growth cone length (Clarke et al., 2020a). The global sqrt(2^nd^ moment) was used for all subsequent quantitative analyses. For purposes of representation of positions on the axon (Fig 2E and 3C, D), it was found useful to indicate the partial sqrt(2^nd^ moment) in the leading and trailing directions, but note that the global sqrt(2^nd^ moment) does not, in general, equal the sum of the leading and trailing partial moments. Motion of the actin peak position was calculated between successive time points. For assessing correlation of actin peak motion during an interframe interval to static features of the axon, comparison was made to the static value at the start of that interval.

### Statistical Methods

Repeated measures ANOVA was used to assess statistical significance of genotype comparisons. A linear model was generated, with first-order autoregression used as the covariance structure to account for repeated measures from each single cell. Box-Cox transformation was applied to outcome variables with non-normal distribution, using the Shapiro-Wilk test to assess normality of model residuals. Tukey’s method was used to correct for multiple comparisons between the three genotypes. Where other statistical tests were applied they are specified in the text and figure legends (GraphPad Prism, Version 9; San Diego, CA).

Pairwise analysis of parameter correlations was quantified by Kendall tau, with significance assessed by Benjamini-Hochberg FDR. Correlations were considered significant at FDR < 5% after correction for multiple testing

PCA was performed by applying standard methods of principal component regression to the wild type dataset. Data from the *UAS-FP4-mito* and *UAS-ena* datasets were then visualized by projecting them into the wild type parameter space.

### Wavelet analysis

The Daubechies type 4 (D4) wavelet transform, modified to account for distribution boundaries, was applied as described previously (Clarke et al., 2020b) to quantify spatial frequency components of the actin distribution:

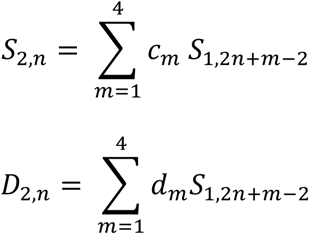

where *N_B_* = the binned actin intensity profile, n = 1, 2, … *N_B_* /4, *c_m_* specifies the low pass filter components, and *d_m_* specifies the high pass filter components. Intensity data were binned such that N_B_ = 2048 with bin width= 0.06μm. Bins overlapping the ends of the distribution were padded with 0 to avoid edge artifacts.

### Jensen-Shannon Divergence (JSD)

For each cell, Jensen-Shannon divergence was calculated between the starting actin distribution and the distribution in each subsequent time point for that cell by the formula

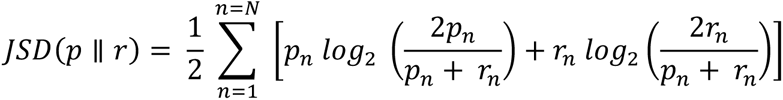

where p and r are the two actin distributions. For three cells expressing *UAS-FP4-mito* (FP4-mito cell #1, #6 and #12) the absolute intensity of the LifeAct signal was quite low (nominal integrated actin intensity per time point < 5 x10^4^ arbitrary units), causing the resulting actin distributions to be discontinuous. Such distributions were not appropriate for JSD analysis and were excluded from this calculation.

### Reproducibility and data exclusion

Movies were not collected or analyzed from cells that failed to show dynamics upon mounting, axons that grew out of the field of focus, or those for which image intensity was too low to detect. One cell (FP4-mito#7) had robust CD4-td-Tomato signal, but insufficient LifeAct-GFP intensity to segment in Imaris and MIPAV. Therefore, this cell was included in analysis of morphological features but not actin parameters. Sample randomization and blinding were not relevant to the experimental design. Sample size was selected based on previous experience with this study design.

## Data and code availability

Numerical data for all figures are included in Supplemental Datasheet. MIPAV code, including plug-ins, is freely available on the NIH website. Python scripts and all other primary data will be deposited in a public depository upon publication.

## Acknowledgements

We are grateful to all the members of our labs for their extensive contributions to the conception, execution, and interpretation of these experiments, particularly Ginger Hunter, Ram Kannan, and Arvind Shukla. We are also deeply grateful to Lenny Campanello for insightful critiques and suggestions concerning our analytical methods, Tianxia Wu for expert assistance with statistical analysis of our time course data, Camille Hanes for assisting in collection and analysis of endstage wing samples for determining terminal phenotypes of *ena*, and Joy Gu and Irina Kuzina for outstanding technical assistance throughout the course of these studies. We thank Holly Cline and David Miller III for their thoughtful comments on the manuscript. We are also indebted to Julie Gates for providing the flies bearing *UAS-ena* and *UAS-FP4-mito-eGFP* transgenes without which these experiments could not have been performed. Spinning disc microscopy was performed in the Cytogenetics and Microscopy Core Facility of the National Human Genome Research Institute, with the invaluable assistance of Stephen Wincovitch. Drosophila stocks obtained from the Bloomington Drosophila Stock Center (NIH P4OOD018537) were essential to these studies. This work was supported by an MURI grant to WG (AFOSR grant number FA9550-16-1-0052), National Science Foundation grants CHE-1800418 and CHE-2102684 to GAP, and by funds from the Basic Neuroscience Program of the Intramural Research Program of the National Institute of Neurological Disorders and Stroke of the National Institutes of Health (Z01-NS003013, to EG).

## Legends to Supplemental Figures

**Supplemental Figure 1.**
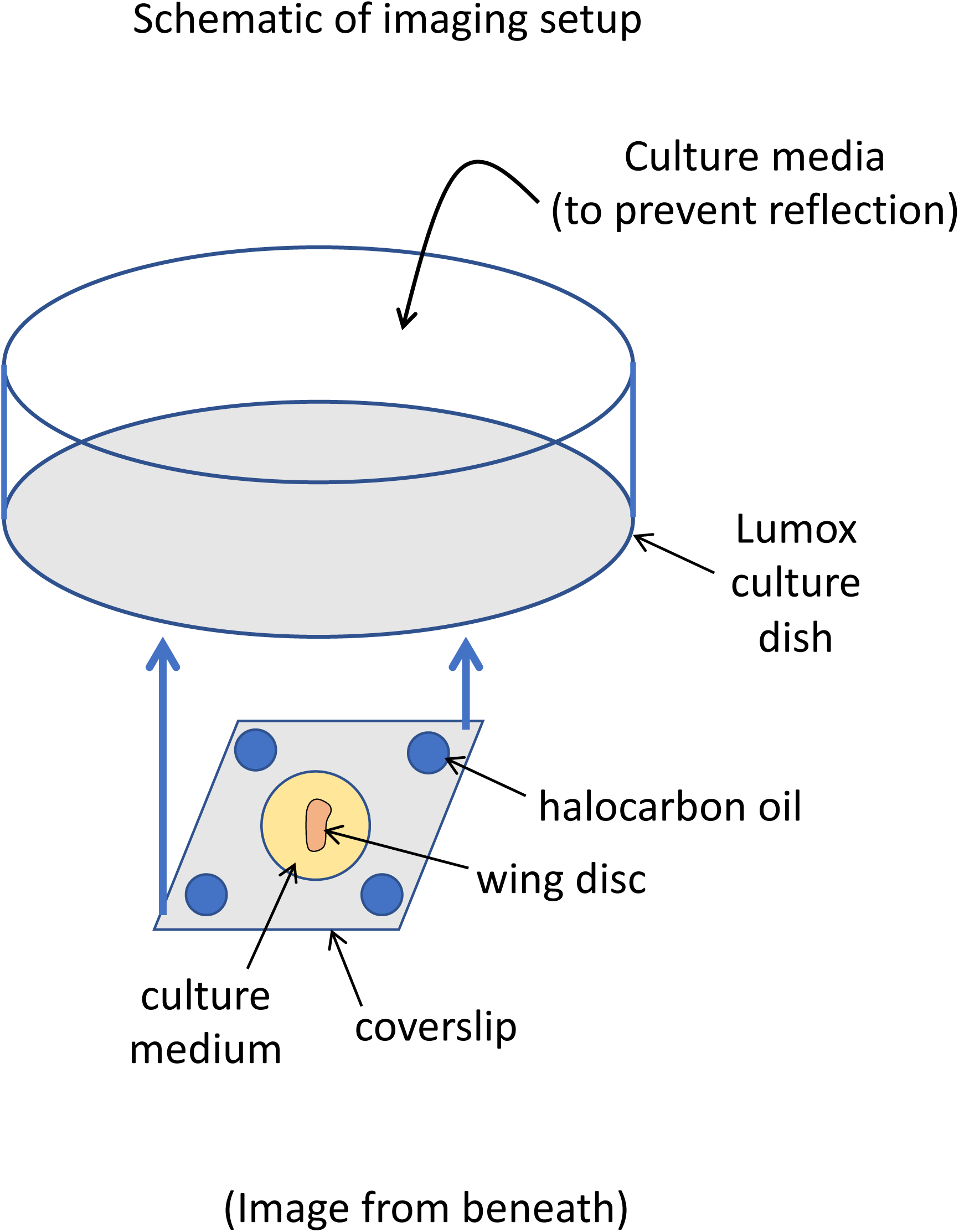
Schematic of imaging mount. Wing discs were mounted in a drop of Schneider’s medium with 10% serum on a #1.5 coverslip and sealed with halocarbon oil to the underside of an air-permeable 35mm Lumox dish. The dish was filled with medium (to minimize reflection) and imaged by spinning disc confocal microscopy on an inverted microscope. See Methods for further details of the mounting method.

**Supplemental Figure 2.**
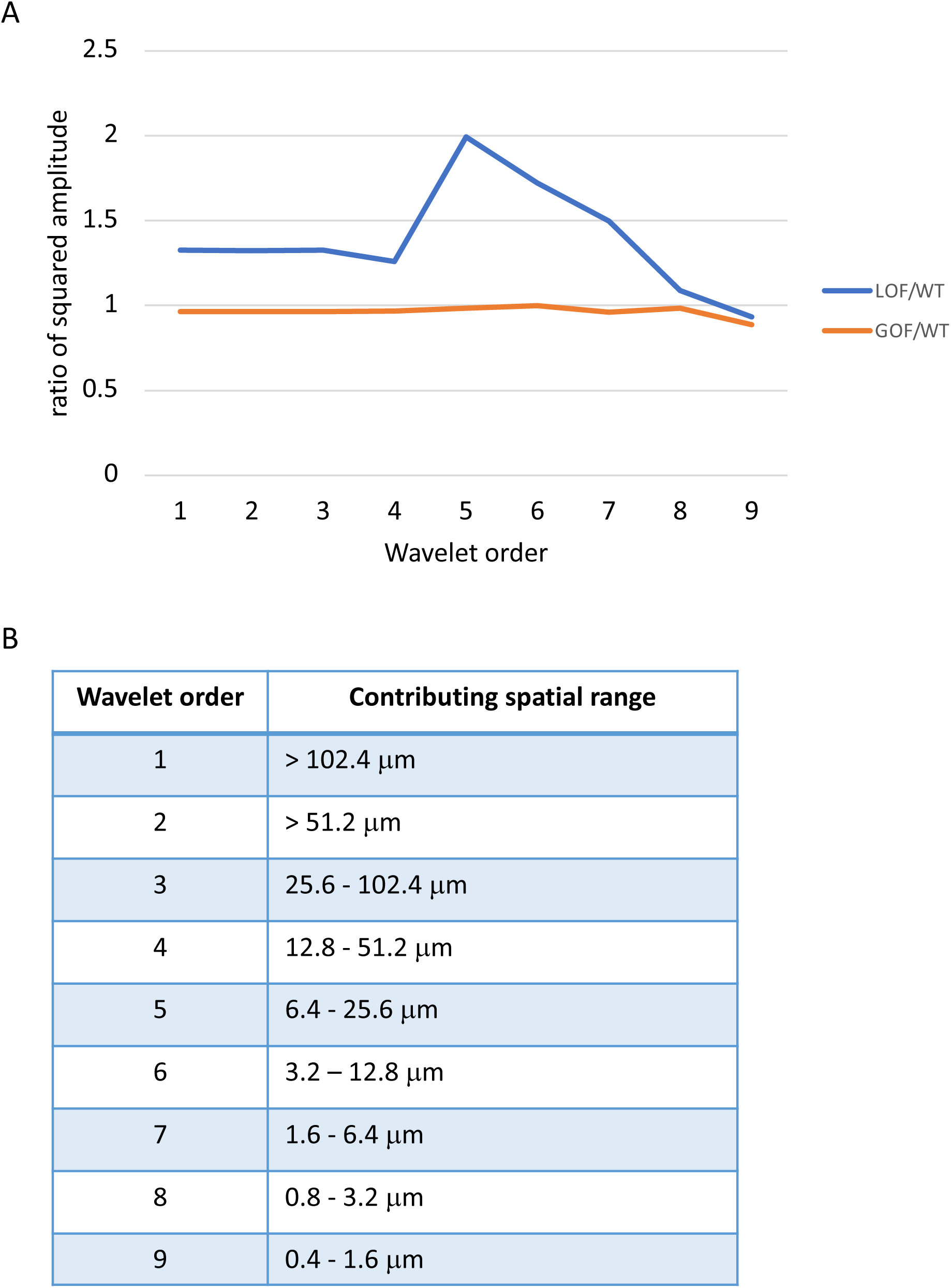
Wavelet coefficients for all genotypes. A. Average (amplitude)^2^ was calculated for each wavelet order from all datapoints for each genotype. Plotted here are the ratio of the (amplitudes)^2^ for *UAS-FP4-mito*/wild type (blue) and *UAS-ena*/wild type (orange). B. Table listing the spatial range corresponding to each wavelet order in this dataset

**Supplemental Figure 3.**
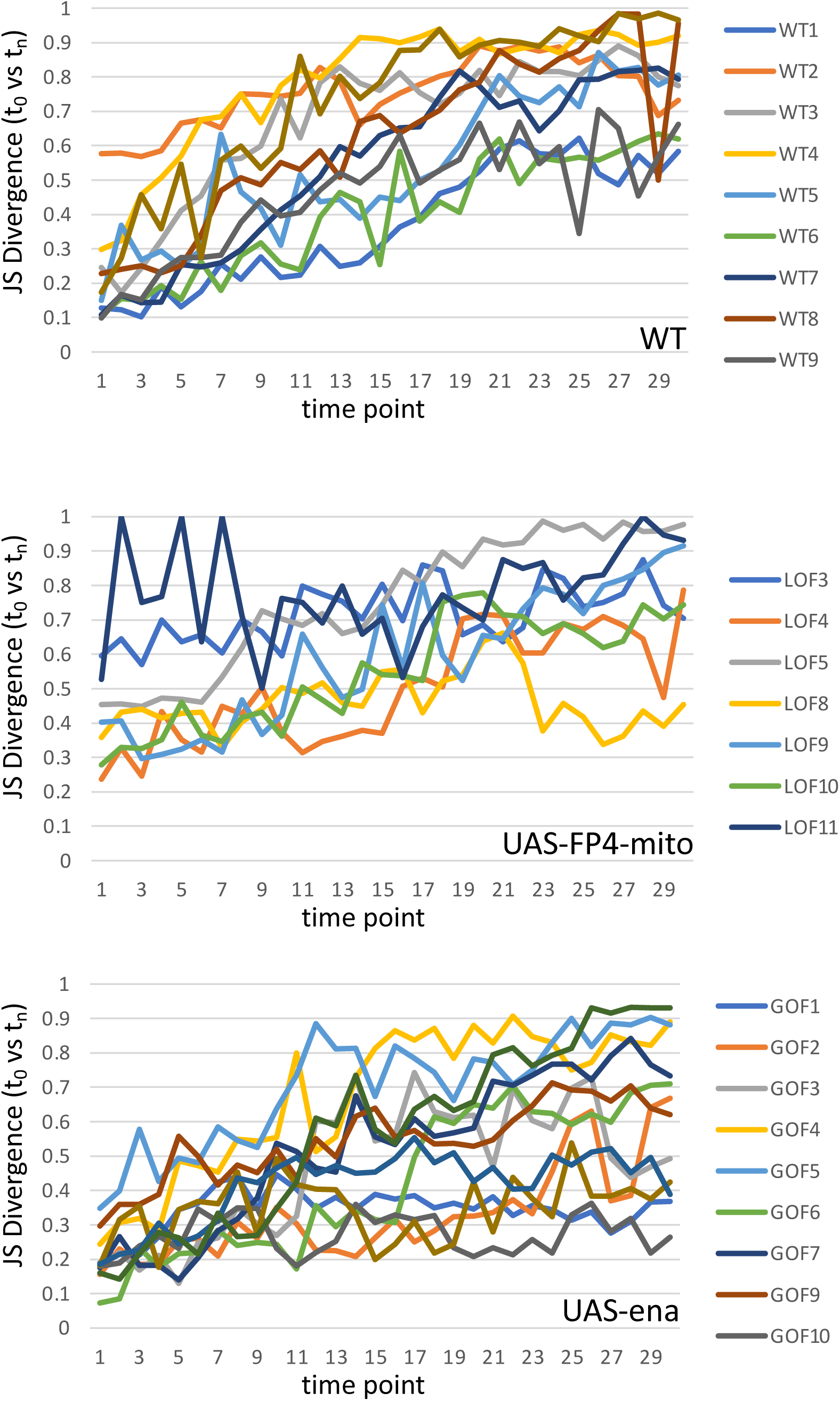
Jensen-Shannon divergence of actin distribution of initial time point vs subsequent time points for all trajectories. Plot of the Jensen-Shannon divergence, calculated between the actin distribution of the starting point of each trajectory and every subsequent time point of that trajectory, vs time. Divergence can vary between 0 (identical distributions) and 1 (unrelated distributions). Time points are numbered on the x-axis, and legend shows the color code identifying the imaged cell. Note that low absolute intensity of the LifeAct signal in three cells expressing FP4-mito caused the measured actin distribution to be discontinuous, and therefore inappropriate for calculation of JSD. Those cells were excluded from this analysis.

**Supplemental Figure 4.**
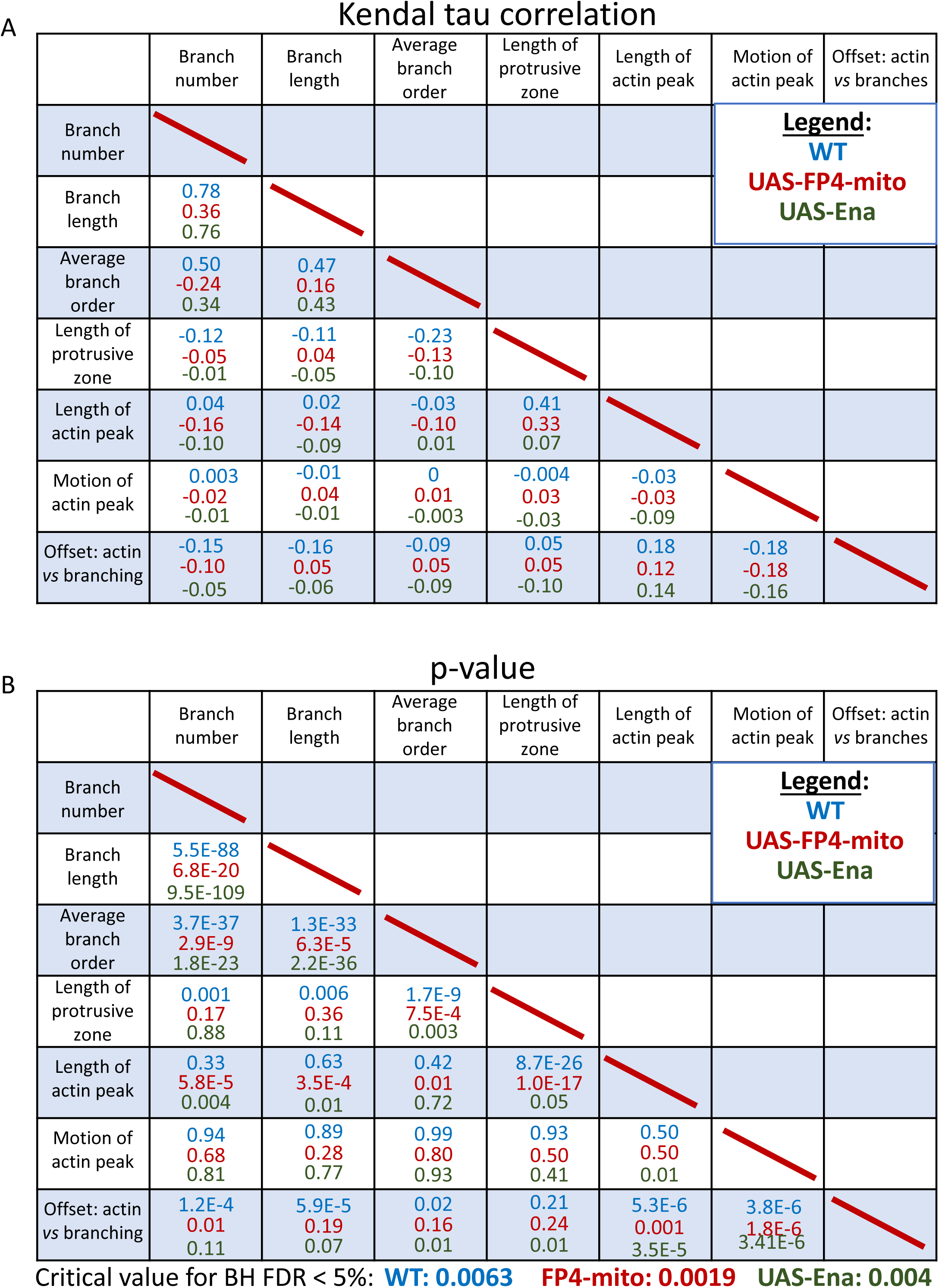
Pairwise correlations of all measured parameters for all genotypes. Kendall tau correlation was calculated for all pairwise combinations of the indicated parameters for every timepoint of each cell imaged, and significance assessed by the Benjamini-Hochberg method. Data for wild type is shown in blue, *UAS-FP4-mito* in red, and *UAS-ena* in green. A) Correlation value: Kendall tau B) Statistical significance: p-value; critical value for FDR < 5% for each genotype (corrected for multiple testing) is indicated at the bottom.

**Supplemental Figure 5.**
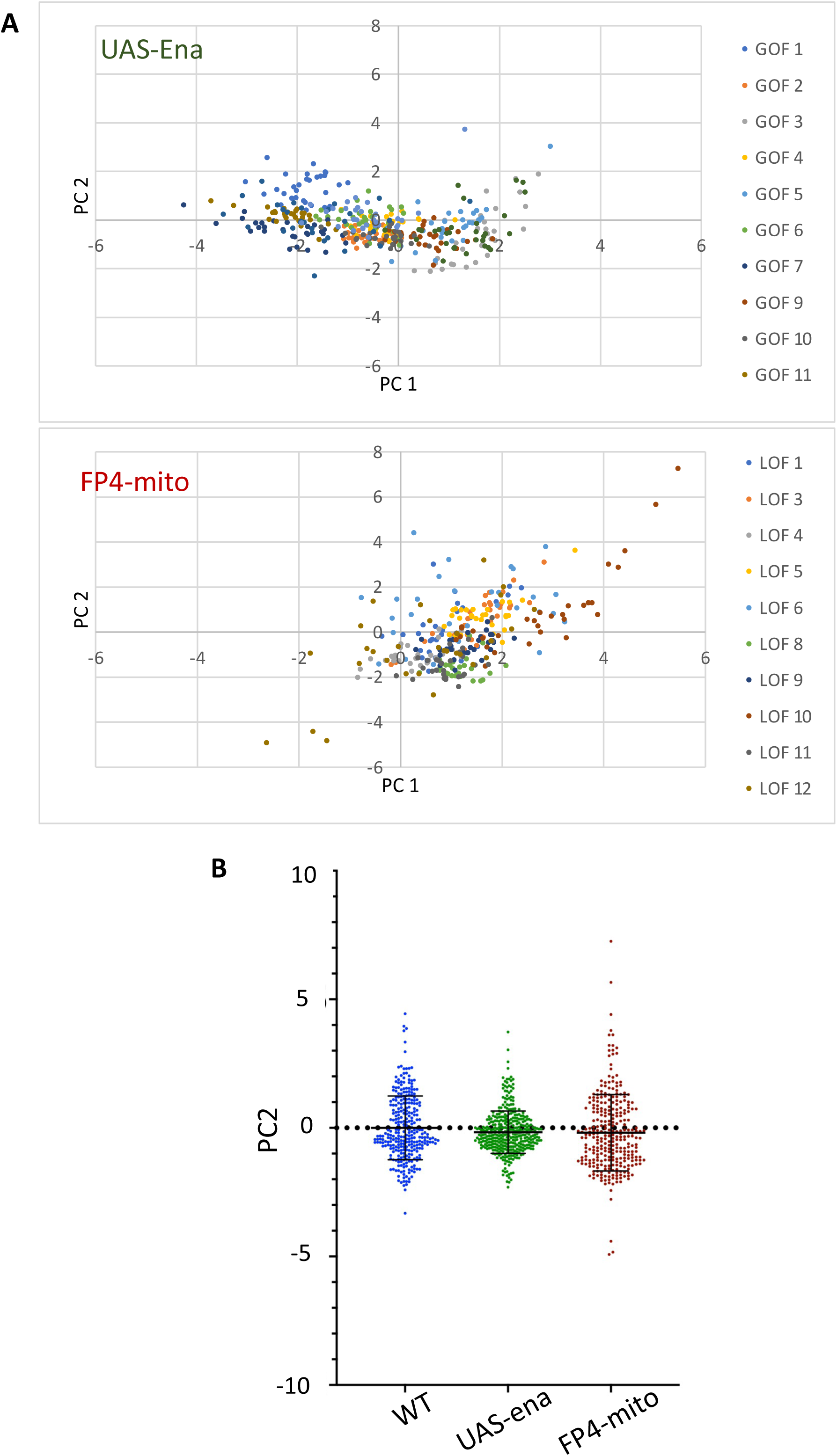
PCA of growth cone parameters, coded by cell and by genotype. A. PCA is shown for *UAS-Ena* and *UAS-FP4-mito* time points, coded by trajectory. Note that UAS-Ena trajectories fail to split into distinct classes PC1 < and > -0.5. B. PC2 values were tabulated for all datapoints of each genotype. Mean and SD are indicated.

